# Automated segmentation of soft X-ray tomography: native cellular structure with sub-micron resolution at high throughput for whole-cell quantitative imaging in yeast

**DOI:** 10.1101/2024.10.31.621371

**Authors:** Jianhua Chen, Mary Mirvis, Axel Ekman, Bieke Vanslembrouck, Mark Le Gros, Carolyn Larabell, Wallace F. Marshall

## Abstract

Soft X-ray tomography (SXT) is an invaluable tool for quantitatively analyzing cellular structures at sub-optical isotropic resolution. However, it has traditionally depended on manual segmentation, limiting its scalability for large datasets. Here, we leverage a deep learning-based auto-segmentation pipeline to segment and label cellular structures in hundreds of cells across three *Saccharomyces cerevisiae* strains. This task-based pipeline employs manual iterative refinement to improve segmentation accuracy for key structures, including the cell body, nucleus, vacuole, and lipid droplets, enabling high-throughput and precise phenotypic analysis. Using this approach, we quantitatively compared the 3D whole-cell morphometric characteristics of wild-type, VPH1-GFP, and *vac14* strains, uncovering detailed strain-specific cell and organelle size and shape variations. We show the utility of SXT data for precise 3D curvature analysis of entire organelles and cells and detection of fine morphological features using surface meshes. Our approach facilitates comparative analyses with high spatial precision and statistical throughput, uncovering subtle morphological features at the single cell and population level. This workflow significantly enhances our ability to characterize cell anatomy and supports scalable studies on the mesoscale, with applications in investigating cellular architecture, organelle biology, and genetic research across diverse biological contexts.

**Significance Statement:**

- Soft X-ray tomography offers many powerful features for whole-cell multi-organelle imaging, but, like other high resolution volumetric imaging modalities, is typically limited by low throughput due to laborious segmentation.
- Auto-segmentation for soft X-ray tomography overcomes this limitation, enabling statistical 3D morphometric analysis of multiple organelles in whole cells across cell populations.
- The combination of high 3D resolution of SXT data with statistically useful throughput represents an avenue for more thorough characterizations of cells *in toto* and opens new mesoscale biological questions and statistical whole-cell modeling of organelle and cell morphology, interactions, and responses to perturbations.

## Introduction

The cellular mesoscale, spanning large molecular complexes to the largest subcellular structures including organelles and cell-wide cytoskeletal assemblies, is emerging as a new frontier in cell biology (A. A. Ekman et al., 2017; Goodsell et al., 2020; Sear et al., 2015). At this scale of biological organization, bulk molecular activities culminate with bulk biophysical effects to produce highly complex cell-wide functions such as cell division, organelle trafficking, and metabolism coordinated across multiple compartments (Qian & Beltran, 2022). The immense complexity of the mesoscale requires novel tools, technologies, and analytical approaches to yield insights into how structure and function are coordinated at the whole-cell level.

The detailed study of intricate cellular structures through microscopy plays a central role in understanding the complex mechanisms governing cellular function and pathology. The budding yeast *Saccharomyces cerevisiae*, is an indispensable model organism in understanding molecular pathways involved in organelle size and shape determination, owing to its powerful genetic tractability and similarity to multicellular animals, including humans, combined with its amenability to imaging methods such as thin-section electron microscopy. *S. cerevisiae* has long been used as a primary model for studying myriad fundamental cellular processes, including, in recent years, the advent of the field of organelle-organelle interactions (Elbaz-Alon et al., 2014; Hariri et al., 2018; Hughes & Gottschling, 2012; Kornmann et al., 2009; Murley et al., 2013; Shai et al., 2018, and many others). However, the small size of yeast cells and their organelles have made it a challenging system for light microscopy-based analysis of organelles. As a result, much of what is known about yeast cell architecture has come from thin-section transmission electron microscopy, a method that gives high resolution but at the cost of extremely tedious sample preparation, which effectively limits studies to either random sections or else serial sections of a very small number of cells. Confocal fluorescence microscopy has been extensively used to image yeast cells and organelles, and by serving as the foundation for integration with other super-resolution imaging modalities, excellent spatial and temporal resolution have been achieved. However, with all such confocal methods, the spatial resolution is anisotropic due to the asymmetric nature of the point spread function (PSF).

In contrast to the low throughput of thin-section EM, and the anisotropic spatial resolution of confocal microscopy, soft X-ray tomography emerges as a powerful option, offering unparalleled advantages for the non-invasive and comparatively high throughput imaging of cells. Unlike other methods, SXT does not require the introduction of external markers, therefore preserving the native state of the cell. This imaging modality, which uses low-energy x-rays, generates high-contrast 3D images of the cells based on the differential X-ray absorption properties of subcellular components. Since the absorption is linear, quantitative linear absorption coefficient (LAC) values are generated that reflect compositional and density variations of structures at the single-cell level. By using full-rotation tomography, the resulting images are completely isotropic in resolution. These unique capabilities are exceptionally valuable for quantitatively assessing the morphological and structural changes in yeast cells, thereby differentiating their phenotypic alterations. Importantly, multiple yeast cells can be imaged in their entirety in less than 10 minutes (Do et al., 2015; Le Gros et al., 2014; Parkinson DY et al., 2008). The ability to image entire cells is important because it allows the full organelle complement of each cell to be visualized and quantified, along with information about the overall cell volume and surface that would be impossible to determine from small sub-images of a cell typical of other high-resolution methods like cryo-EM.

The ability to rapidly collect whole-cell images from large numbers of cells in a single experiment leads to a new challenge - how to identify the individual organelles from each label-free 3D image dataset. This process of data segmentation, which enables researchers to isolate specific organelles and examine the structural parameters within the cell, is of critical importance for accurate segmentation in yeast cell biology, as it underpins all further analysis and interpretation of cellular morphology. It is particularly fundamental to any efforts to use SXT as a routine method for quantifying cell anatomy, defined as the cumulative morphology and positioning of several organelles in the whole cell context. The ability to accurately segment and analyze images of cells and their components observed from various modalities facilitates a deeper understanding of cellular functions, behaviors, and interactions (Loconte et al., 2022, 2023). For high-resolution contrast-based microscopy modalities, especially volumetric modalities such as SXT, segmentation is typically done manually. This is an extremely laborious and time-intensive task, requiring hours, or in some cases, days, to segment a single cell, depending on the number of structures being segmented per image. Automating this bottleneck step of manual image segmentation allows for the rapid processing of large volumes of data. Auto-segmentation leverages computational algorithms to dissect images into their fundamental components, delineating these segments based on their distinctive features and properties. This process facilitates the precise identification and quantification of subcellular structures in a high-throughput manner. This is particularly indispensable for current large-scale data-intensive approaches in cell science, in which analyses of large datasets are increasingly required to draw meaningful conclusions.

The advancement of deep learning-based methods has shown great potential in achieving efficient and accurate auto-segmentation (Dyhr et al., 2023; Egebjerg et al., 2023; A. Ekman et al., 2020; Erozan et al., 2024; Nahas et al., 2022; Pelt & Sethian, 2018). Convolutional neural networks (CNNs), especially U-net in its various forms have proven highly effective in tasks such as medical image segmentation and object recognition (Azad et al., 2024; Fu et al., 2021; Siddique et al., 2021), where their hierarchical and learned representations enable accurate and robust delineation of regions of interest. CNN offers a versatile, functional space that encompasses both the extracted features and the combinations with a single trainable optimization function.

In this work, we combine soft X-ray imaging, deep learning auto-segmentation, genetic manipulation of yeast cells, and a robust method for analyzing and quantifying subcellular structures in whole cells, resulting in a workflow for large-scale analysis of cell anatomy at the whole-cell level. We describe the full process and technical considerations of developing a custom auto-segmentation algorithm and demonstrate the potential for statistical analysis of organelle and cell morphometrics and biophysical properties across genetic conditions and detailed global and local shape analysis using meshing of isotropic volumes. The integration of quantitative analysis with high-resolution imaging aims to bridge the gap between cell biology and genetics, providing a holistic view of cellular anatomy and its functional implications potentially paving the way for advancements in genetic engineering, therapeutic development, and beyond.

## Results

### Comparing whole-cell reconstructions of budding yeast from confocal fluorescence imaging and soft X-ray tomography

We generated 3D soft X-ray tomographic reconstructions of cryo-preserved log-phase yeast cells based on organelle segmentations. Fig. 1A shows the 3D surface rendering of multiple budding yeast cells at various stages of the cell cycle within a 12x12x12 µm field of view. This high-throughput imaging modality, relative to modalities of comparable resolution such as serial sectioning EM, with a data acquisition time of 5 minutes for 8 yeast cells, non-destructively captures both the external contours of the entire cell and the internal organelle structures without the need for additional labeling, thereby revealing the morphologies in their native state. To quantify and analyze the biophysical properties of the cell structures, we then manually segmented the subcellular organelles based on their distinct morphologies and linear absorption coefficients (LACs). Fig. 1B demonstrates the manual segmentation process of organelles in a dividing yeast cell, underscoring the ability of SXT to provide high-resolution, quantitative data that can be used to analyze morphology and compositions accurately. Fig. 1C depicts the LAC distribution across different organelles. The mean LAC, along with their standard deviations (S.D.), are provided for vacuoles, nuclei, lipid droplets, mitochondria, and cytosol, which are 0.182 μm^-1^ (±0.039 μm^-1^), 0.310 μm^-1^ (±0.020 μm^-1^), 0.570 μm^-1^ (±0.111 μm^-1^), 0.382 μm^-1^ (±0.018 μm^-1^), and 0.346 μm^-1^ (±0.027 μm^-1^) respectively. These measurements allow for the differentiation of organelle compositional density and 3D spatial distribution.

**Fig 1.**
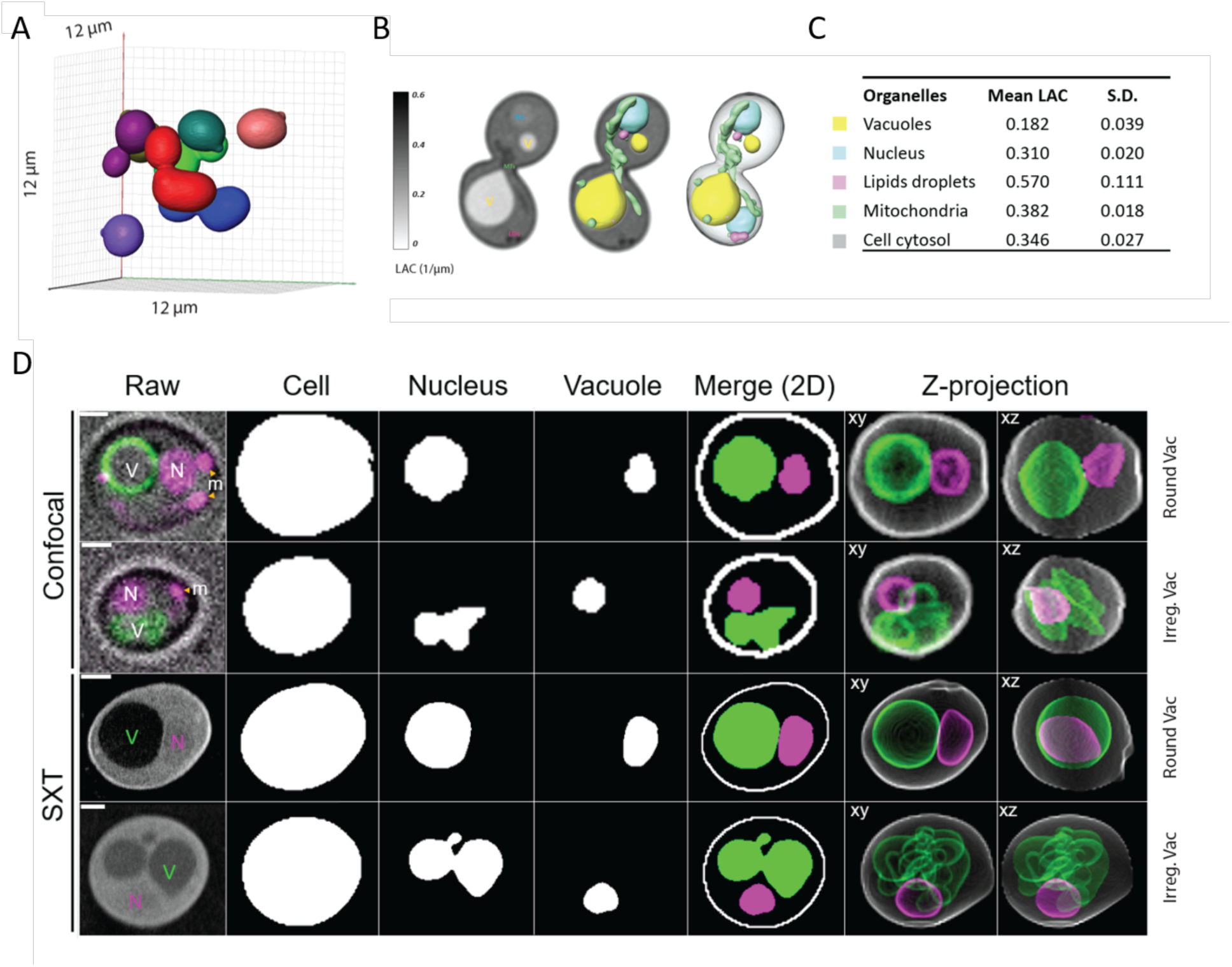
SXT imaging and segmentation of yeast cells. A) 3D rendering of yeast cells in a single tomographic reconstruction. B) Manual segmentation of organelles in single cell tomograms based on LAC. C) Mean and standard deviations of LACs of segmented organelles. D) Examples of single yeast cell 3D raw data and segmentations. (i, ii) Confocal micrographs with vacuole (V) marked by VPH1-GFP and nucleus (N) marked by DAPI. DAPI also stains mitochondrial DNA (m). (iii, iv) SXT raw tomograms. Segmented cell, nucleus, and vacuole are shown as representative 2D slices and sum slices z-projections of edge detections of segmentations. 3D rotations of each cell are provided in Supplementary Movies 1-4.

As a proof of concept of the quality of image data and cellular information achievable in whole-cell SXT imaging, we show a side-by-side comparison of single-cell examples from standard confocal microscopy and SXT, including segmentations and 3D reconstructions of the cell, vacuole, and nuclear volumes for each cell (Fig. 1D and Supplementary Movies 1-4). Two representative cells from each modality, one with a spherical vacuole and one with a vacuole of irregular geometry, are displayed. The confocal data have a near-isotropic voxel size of 93 nm laterally (xy) and 100 nm axially (z), and the SXT data have an isotropic voxel size of 30-36 nm, with exact resolution varying slightly between individual tomograms across our dataset. SXT data were initially manually segmented based on the LAC, reflected in the visible contrast among structures in the raw tomogram. SXT has clear advantages for generating more realistic 3D representations of cells compared to confocal, providing more precise object boundaries for morphometric quantification, and revealing detailed shape information in complex structures such as clustered or irregularly shaped vacuoles. While confocal deconvolution improves the distortion of optical data in the z-direction, SXT is nearly isotropic and does not show any distortion along any axis. The label-free nature of SXT of cells circumvents concerns of a non-specific signal, the perturbative potential of fluorescent labels and phototoxicity, as well as possible shape distortions arising from organelle movements during live cell image acquisition. Thus, SXT offers superior 3D structural resolution, with multi-organelle information at the whole cell level. However, the accessible structural information at this resolution lags behind optical modalities due to the limited throughput of manual segmentation.

### The application of machine learning and convolutional neural networks to segmentation

To date, most studies that quantitatively analyze cellular structures at sub-optical resolution still heavily rely on manual labeling – a time-consuming, labor-intensive process that often depends on the individual’s expertise in identifying features within the reconstruction. As we move towards high-throughput analysis, manual labeling becomes increasingly impractical due to the growing number of samples. With the emergence of promising automatic segmentation methods (Çiçek et al., 2016; Pelt & Sethian, 2018), it is crucial to develop workflows that effectively integrate these tools into the structural analysis pipeline for large datasets. To meet this need, we have employed a task-based segmentation pipeline. Each research question requires a specific set of semantic needs. We aim to create a pipeline that can collect the necessary data, choose an appropriate model architecture, and train a task-specific algorithm tailored to the user’s specific inquiries. Here, we present an example of a fully 3D U-Net-type Convolutional Neural Network (CNN).

In the case of this work, the data were raw 3D SXT images of yeast cells, and the task was to use ML to aid in the segmentation of the individual cells and their semantic labeling into cell, nucleus, vacuole, and lipid droplets. The outline of the iterative process is shown in Figure 2. We start with an initial set of manually labeled data (n=7), from which an initial model is trained. This model is then used to generate automatic labels for the whole set of raw data from hundreds of cells. From this set of initial results, the user then selects poor-quality segmentations and either manually segments or refines them to be used as additional training data for the next iteration. The cells were first pre-segmented, allowing each individual cell to be segmented separately. This instance segmentation as a preprocessing step significantly accelerates the training process, as the training data doesn’t contain “empty” data sets, and the problem with partial cells (cells clipped by the boundary of the image) that can be difficult to label manually is avoided. The pre-segmentation involved identifying the cells within the capillary and using watershed segmentation for separation, and allowed for the use of partially labeled data (raw tomograms, where only one cell was manually labeled). The progression of this partially manual segmentation process with increasing numbers of cells is shown in Figure 2B. A key challenge lies in identifying the optimal stopping point. With too few samples, the generalization of the model to the dataset is poor. Conversely, excessively continuing the iterative process diminishes the method’s advantage, essentially converging towards manually segmenting the entire dataset.

**Fig 2.**
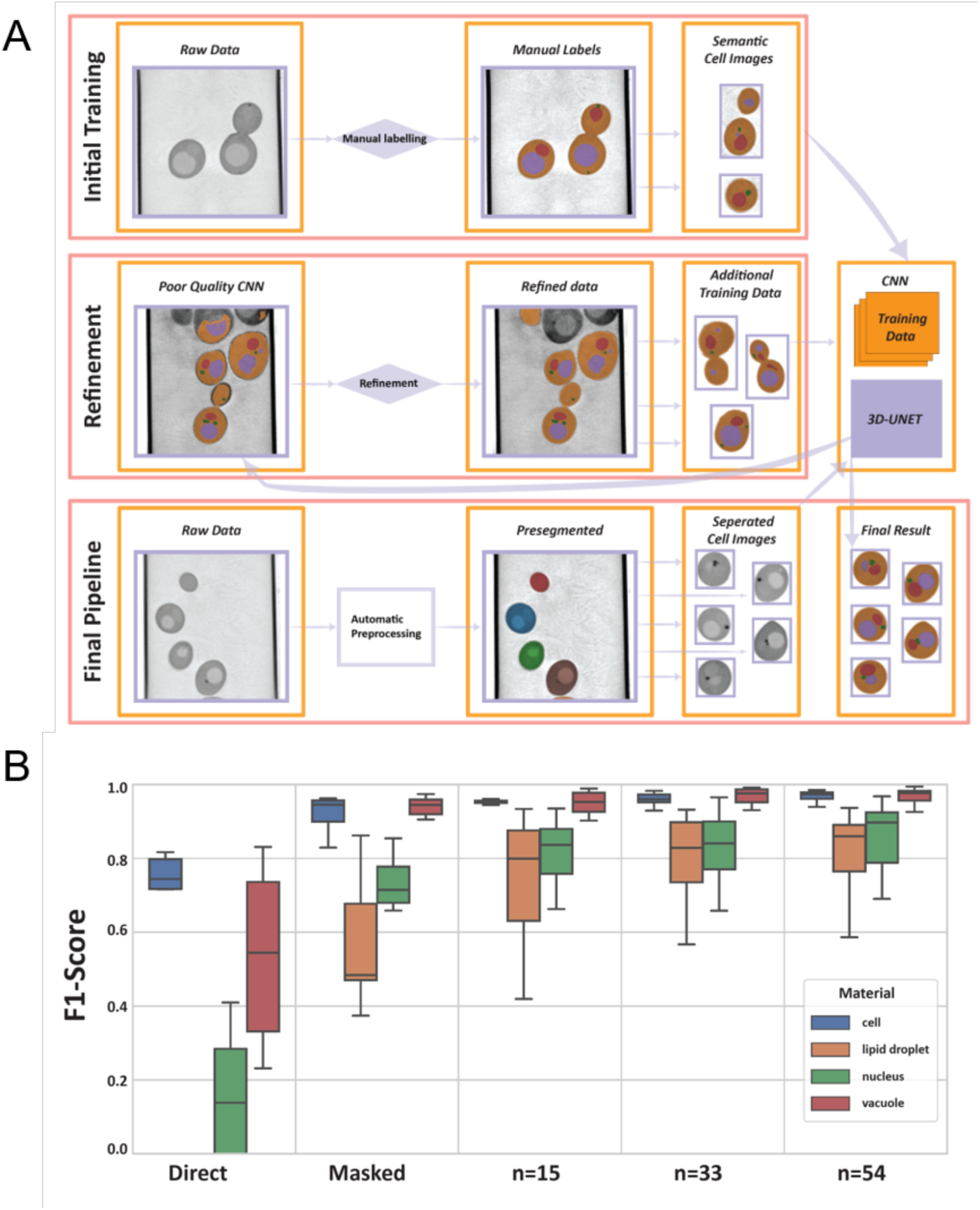
Overview of the iterative process of segmenting a batch of cell images. A) An initial set of manual labels are used to train the model. Poor quality inferences from the initial model were refined through manual correction and added to the training data. This iterative refinement continues until the model achieves satisfactory segmentation quality. B) Improvement of F1-Scores with manual iterative refinement showing the F1-scores for the different labels (cell, lipids, nucleus, vacuole). The “Direct” column shows the initial F1-scores trained using full volume. The “Masked” column and subsequent columns show results of using pre-segmentation of the individual cells, prior to training.

The metrics used to train the segmentation models do not directly address the usability of results in terms of the research question. Furthermore, the required loss metric of segmentation to answer research questions depends on the metrics that one wants to extract. Erozan, *et al*. (2024) showed that a direct U-Net is well-suited for segmenting and assessing cell volume but can produce statistically significant differences in metrics such as surface area when compared to the training data, indicating the need for additional refinement or alternative methods. Volumetric measurements, such as total volume, mean LAC, or structural thickness, may not require as precise segmentations at the boundary, whereas surface area measurements and minimum distance-based metrics (like structural thickness) are highly sensitive to single voxel errors, which are not well captured by voxel-based loss functions.

In Fig. 3A, we show the distribution of the label field volumes in both the training data and model predictions. Although the F1 score for the nucleus label is substantially lower than that of the vacuole, the automatic segmentation does not produce significantly different results in terms of volume when assessing population statistics. Conversely, for lipid droplet volumes in the case of VPH1-GFP, the automatic segmentation results show significant deviations from the manual labels, indicating that these results require further evaluation for reliability. Fig. 3B provides examples where CNN has overestimated or underestimated the volume of lipid droplets (LD) compared to the training data, highlighting the need for careful evaluation and potential refinement of the automatic segmentation results. Overall, the results show robust automated segmentation across all strains, particularly for the nucleus and vacuole, enabling quantitative comparative analysis of cell lines, only requiring a fraction of the collected data to be manually segmented.

**Fig 3.**
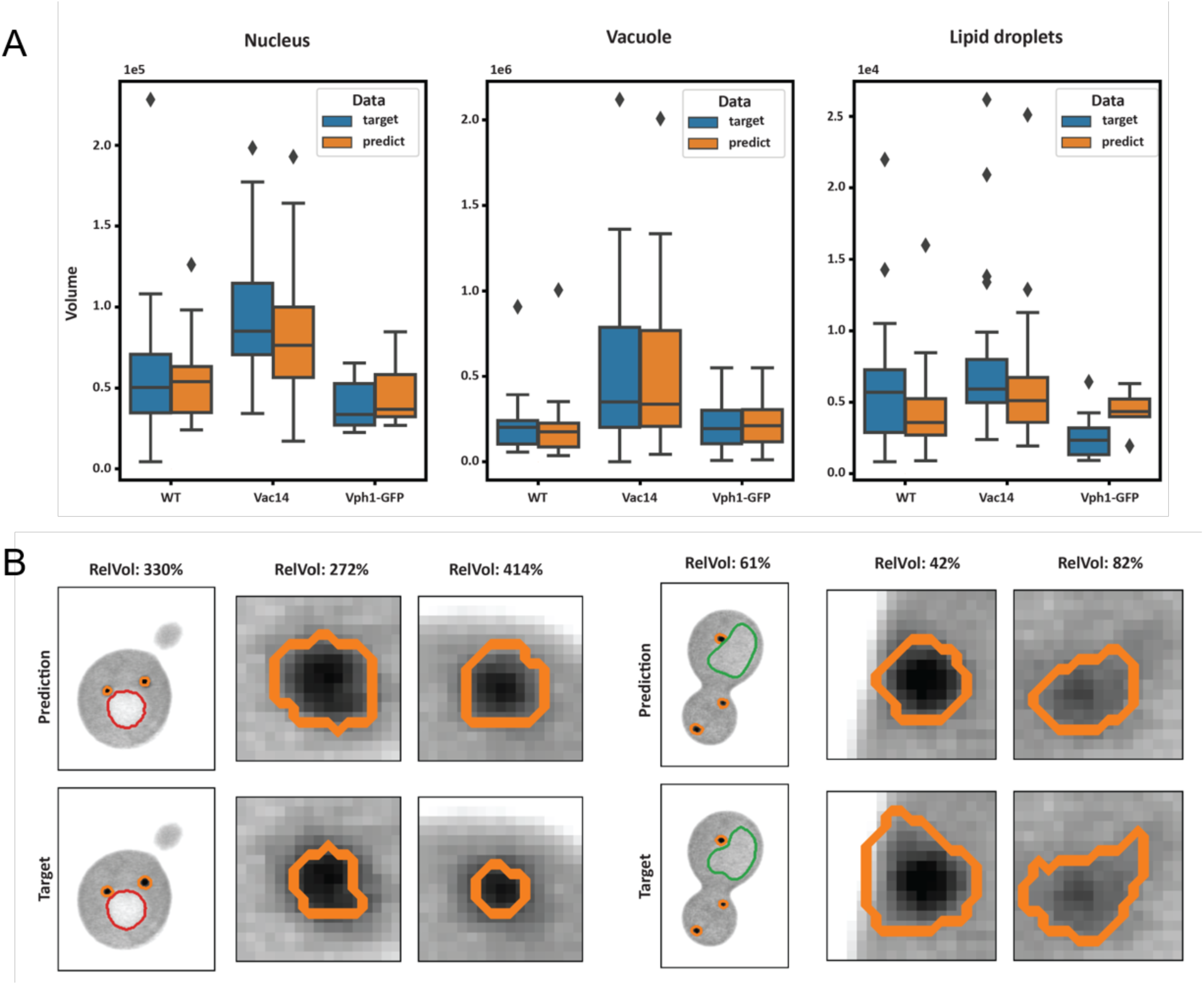
Validation of auto-segmentation. A) The volume measured from the training data and predictions, respectively for the three different identified organelles. The volume predictions for the nucleus and vacuole do not deviate significantly from the manual labels, indicating that the model can reliably be used to describe population statistics of these cell structures. However, the segmentation of lipids shows considerable deviation from the manual data, suggesting that further refinement and quality assessment are needed for accurate lipid volume predictions. B) Example slices of two outliers in the lipid volumes from population VPH1-GFP (left) and WT (right). Zoomed in slices show two examples of the labeled lipid droplets (orange contour). RelVol depicts the relative volume of the predicted lipid label to the (manually segmented) target label. Red and green contours are of nucleus and vacuole, respectively.

### Quantitative phenotypic analysis from SXT data comparing auto-segmented yeast strain morphometrics

We used the auto-segmentation outputs in the form of voxel-based binary image stacks to carry out a statistical 3D morphometric characterization of cells and organelles in unbudded early G1 cells in three related strains: the haploid wild-type strain BY4741 (WT, n=287); a strain expressing VPH1-GFP fusion marking the vacuole membrane in a BY4741 background (Chan et al., 2016; Chan & Marshall, 2014) (VPH1-GFP, n=138); and *vac14d*, a large-vacuole mutant, originally identified as a class III vacuole inheritance mutant with an enlarged and fused vacuole phenotype, involved in PI(3,5)P2 metabolism (Banta et al., 1988; Bonangelino et al., 2002; Rivero-Ríos & Weisman, 2022; Rudge et al., 2004), in the BY4741 VPH1-GFP background, (*vac14*, n=65). These measurements are based on voxel counts, scaled to isotropic voxel sizes in the range of 30-36 nm. Statistical summarization of the sample distributions across strains showed that volume distributions and volume fractions were generally similar to previously reported values for yeast cells, vacuoles (Chan & Marshall, 2014; Uchida et al., 2011), nuclei (Jorgensen et al., 2007), and lipid droplets (Egebjerg et al., 2023; Uchida et al., 2011) (Fig. 4, statistics in Supplementary Table 1). We observed the highest variability in volume and proportional volume in vacuoles, as compared to nuclei and lipid droplets (Fig. 4A-E). Vacuoles and nuclei had sphericity values with a median of 0.64-0.66 in all conditions, but vacuoles showed a higher variance in shape compared with nuclei (Fig. 4FG).

**Fig 4.**
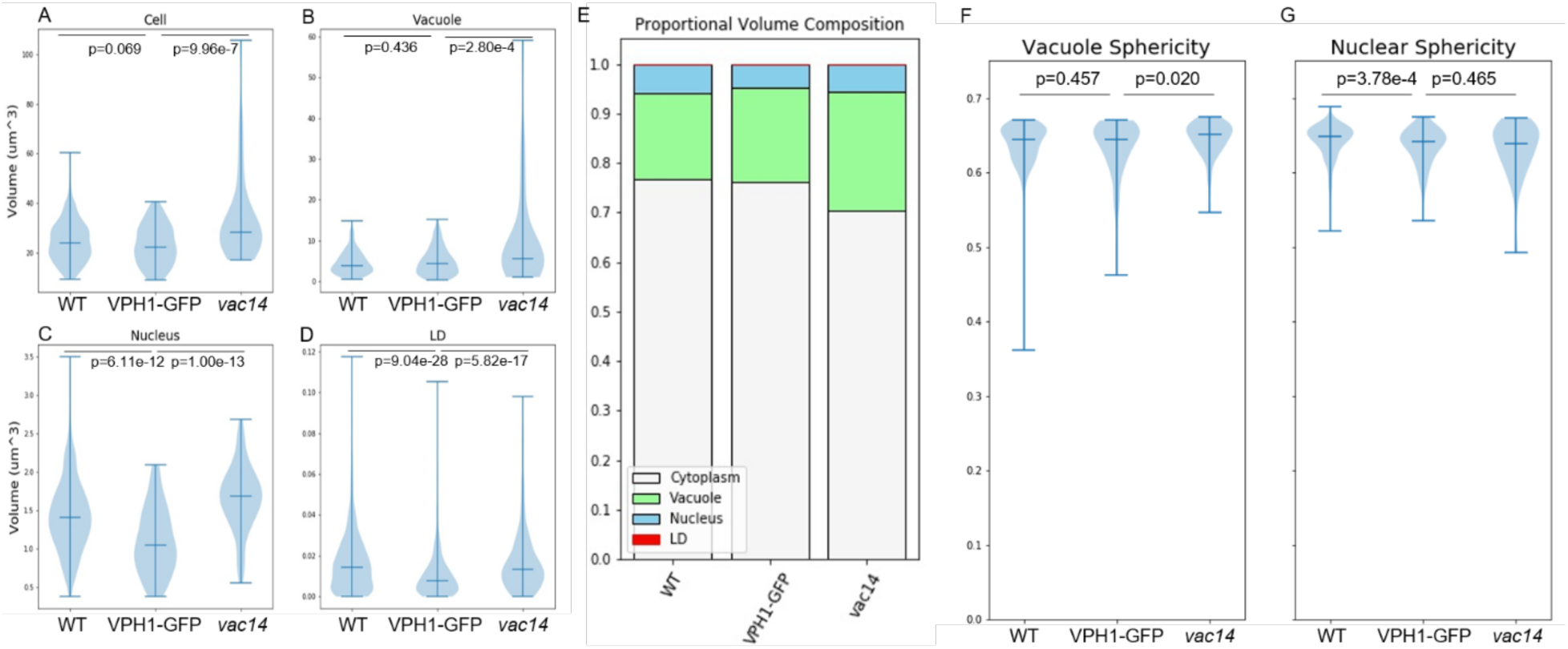
Quantitative phenotypic analysis from SXT data comparing auto-segmented yeast strain morphometrics. Volume distributions for (A) cell, (B) vacuole, (C) nucleus, and (D) lipid droplets plotted as violin plots for WT, VPH1-GFP, and *vac14* strains. Midline marks the median. (E) Proportional volume composition for each strain. (F) Vacuole and (G) nucleus sphericity distributions plotted as violin plots for each strain, with midline marking the median. P-values reported from two-way Mann-Whitney U-tests for nonparametric distributions. N = 287 (WT), 138 (VPH1-GFP), and 65 (*vac14*). Complete summary and significance test statistics reported in Supplementary Table 1.

Fluorescence imaging relies on well-characterized fluorophores with specific localization and no significant impact on cellular structure or function such as viability loss or gross morphological defects. Without a way to know whether a fluorescent tag alters protein function or cellular morphology in less consequential ways, imaging of tagged strains is always associated with the potential risk that the tag itself may subtly affect organelle size or shape, particularly when, as is often the case, the proteins tagged are involved in organelle biogenesis, function, inheritance, or interactions. Due to the label-free nature of SXT, our data enables direct morphometric comparison of the parental strain and the strain expressing VPH1-GFP, a V-ATPase component protein commonly used as vacuole membrane marker for fluorescence imaging (Chan et al., 2016; Chan & Marshall, 2014; Shimasawa et al., 2023), allowing us to assess subtle morphological changes potentially caused by VPH1-GFP expression, despite no reported effects on cellular function. We found that VPH1-GFP expression confers small but statistically significant reductions to nuclear and lipid droplet volume distributions, but not to the volume distributions of the cell and vacuole (Fig. 4A-D, Supplementary Table 1). The overall proportional volume composition was very similar between VPH1-GFP and the parental strain (Fig. 4E, Supp. Table 1). Nuclear sphericity was significantly reduced in the VPH1-GFP strain, but there is no significant difference in vacuole sphericity (Fig 4FG, Supp. Table 1). These results illustrate one important potential application of SXT - providing a label-free control to assess the effect of added fluorescent labels on cell structure, and the importance of characterizing controls for quantitative comparisons due to the potential presence of subtle morphological differences.

The VPH1-GFP strain, in turn, served as the direct control for the *Δvac14* strain, which also expresses VPH1-GFP. *Δvac14* is an example of a classical deletion mutant with a known morphological defect, described as “vacuolar enlargement” (Bonangelino et al., 1997; Dove et al., 2002; Rudge et al., 2004). However, a precise quantitative description of the phenotype, such as the degree and variability of volume increase, has not been reported. Comparing VPH1-GFP and VPH1-GFP *Δvac14* showed that the mutation has only a modest effect on the average volume, but a large effect on volume variability (Fig 4B). Cells, nuclei, and LDs, were also larger in *vac14*, such that distribution (right-tailed variance) was increased for cell volume, but central tendency was increased for nuclear volume and LDs. A comparison of proportional volume composition (stacked plot of means) showed that proportional volume of the vacuole increased by 5.5% (18.9% of the cell volume in VPH1-GFP, 24.4% in *vac14*) while the “remaining cytoplasm” (a per-cell metric defined as the known organelle volume subtracted from the cell volume) was reduced by 6% (76.2% in VPH1-GFP, 70.3% in *vac14*). Volume fractions of the nucleus and LDs increased slightly (nucleus: 4.8% in VPH1-GFP, 5.5% in *vac14*; LD: 0.063% in VPH1-GFP, 0.068% in *vac14*) (Fig 4C). LDs occupy less than 0.1% of the cell volume, so the ability to detect variations within LD morphometrics are a significant strength of our approach, although such variations do not constitute significant volumetric changes for the whole cell. Vacuole sphericity was slightly increased with lower variance in *vac14* relative to VPH1-GFP, but we found no difference in nuclear sphericity. Therefore, the *Δvac14* phenotype goes beyond vacuole enlargement to include vacuole size variability, increased sphericity, and overall cell and organelle volume increase.

### High-resolution shape analysis of SXT data with meshing

Organelle shape is fundamentally linked to function. However, it is difficult to precisely quantify organelle shape in three dimensions due to limitations in throughput and resolution in most volumetric data, particularly in the z-axis. The high resolution and isotropy of our SXT data provides the potential for detailed 3D shape analysis of organelles, but the processing and representation of the segmented data as 3D objects are critical steps for unlocking this potential. There are several ways to reconstruct an organelle surface in 3D, including voxel-based reconstructions (projecting the raw image stack in three dimensions), and meshing (representing the three-dimensional surface as a simplified geometrical, typically triangulated, mesh), as exemplified in whole-cell multi-organelle visualizations for a representative cell of each strain (based on our statistical characterizations described in Figure 4) in Supplementary Figure 1 (see full rotations in Supplementary Movies 5-10). Meshing has several advantages for shape analysis over voxel-based reconstructions. Surface information is extracted and simplified from a solid volume, reducing data size and computational burden of processing and analysis. Meshing allows the conditioning of the surface to remove jagged voxel-related artifacts, as well as customization of information density to the unique surface features of the structure, such as reducing the density of information points (triangles) in smoother regions while increasing it in feature-dense regions (Feng et al., 2012; Lee et al., 2020a).

Artifact removal and mesh surface conditioning also facilitate more accurate 3D morphometrics. Volume measurements taken from FIJI 3D Suite (Ollion et al., 2013; Schindelin et al., 2012) for voxel-based segmentations (as in Fig. 4) and from GAMer2 (Lee et al., 2020b) for meshes, both pre- and post-mesh processing, were typically within 5% of each other (Supplementary Table 2). We note <1% volume difference between pre-and post-processed meshes, demonstrating the volume preservation of our mesh refinement pipeline (Materials & Methods). Organelle volume ratios (vacuole:cell and nucleus:cell) were generally consistent between all measurements. However, voxel-based surface area measurements were significantly higher than those produced by mesh morphometrics, highlighting the unsuitability of voxel representations for surface area measurements due to the artifactual overestimation of surface area by voxel edges. Mesh processing had a larger effect on surface area than on volume measurements because smoothing of sharp edges and voxel terracing effects reduces overall surface area by 2-8%.

For shape quantification, sphericity measurements of voxel reconstructions and pre-processed meshes were generally well within 10%, and consistently increased after mesh processing, as expected due to smoothing, resulting in post-processed sphericity measurements typically ∼5% higher than voxel-based measurements (Supplementary Table 2). Meshing with appropriate processing provides highly detailed surface topographies, with the potential to yield rich shape information and in-depth shape analysis beyond what is achievable with voxel-based reconstructions. To illustrate this, we demonstrate surface curvature mapping using a selection of morphologically variable vacuoles. The number of vacuoles present per cell may also be variable, ranging from 1-10 vacuoles per cell depending on the strain (Chan & Marshall, 2014). Clusters of several vacuoles can appear as a large mass or cluster in fluorescence imaging (Fig. 1D), and in voxel reconstructions, the full detail of local and global surface features can be obscured due to occlusion of voxels and quantization artifacts (Fig. S1). Another strategy for representing surfaces that can avoid problems due to voxel artifacts is surface harmonic fitting (Chan & Marshall, 2014; Marshall et al., 1996), however, this approach is limited to “star-shaped” surfaces, i.e. surfaces for which every outward ray from the centroid intersects the surface at a single point. Complex organelle shapes can violate this constraint. Mesh representation is not constrained in this way and can therefore be used even for complex organelle shapes.

We asked whether meshing enables the calculation of more informative global shape metrics for complex organelles compared to crude measures obtainable from voxel reconstructions, such as sphericity. We compared several examples of vacuoles of increasing sizes and shape complexities (Fig. 6, Table 1). Examples were selected to represent the observed sphericity range calculated from voxel-based segmentations (Fig. 4FG), and volume calculated following mesh processing (Table 1). We chose small (A-C, 1.005-1.443 μm^3^), medium (E-F, 5.523-6.316 μm^3^), and large (G-I, 7.250-9.767 μm^3^) vacuoles of increasing complexity, ranging from the most sphere-like vacuoles in our dataset (ADG, 0.670-0.674), to median-range (BEH, 0.628-0.629), to the least spherical vacuoles in the dataset across all strains (CFI, 0.464-0.497). It is immediately apparent that sphericity is not adequate as a metric to distinguish objects based on shape, as the vacuoles in Figs. 6B and 6E have identical sphericities (0.628) but are visually different, with more pronounced protruding features in 6E.

Curvature, a key feature of surface morphology defined as the magnitude of surface deviation from a tangent plane (Feng et al., 2012), can be calculated across the mesh. Principal curvatures at each vertex (κ_1_ and κ_2_) are represented by combined measures such as mean curvature ((κ_1_+κ_2_)/2) or by Gaussian curvature (κ_1_*κ_2_), visualized as a color intensity gradient across the surface of the mesh. Mean curvature can be conventionally interpreted as positive for convex vertices and negative for concave vertices. Gaussian curvature can be interpreted as positive at ellipsoid or spherical regions (both principal curvatures are positive), neutral at paraboloid or cylindrical regions (k_2_ = 0), and negative at hyperboloid or saddle-shaped regions (Feng et al., 2012; Koenderink & van Doorn, 1992). Full curvature maps for each vacuole are shown in Fig. 6A-I.

In order to quantitatively represent the global shape of each vacuole to distinguish between various shapes of increasing complexity using a single metric, a statistical aggregate (average and standard deviation) of mean (*H*) or Gaussian (*K*) curvature values across the entire mesh is calculated (Fig. 6JK, Table 1). A perfect sphere has a positive average mean curvature and Gaussian curvature, with magnitude increasing with radius, and no variance across the mesh. Increasingly complex shapes are expected to feature increasing curvature variance. As expected, as we compare vacuoles of similar sphericity with increasing volume (ADG, BEH, and CFI), mean *H* and *K* slightly decrease toward zero in central tendency (median) but increase slightly in variance. As we compare vacuoles of similar volume with increasing sphericity (ABC, DEF, GHI), there is no meaningful change in the central tendency of *H* and *K*, but variance increases several-fold, with a dramatic increase in variance for the most complex vacuoles (CFI).

Curvature enables the calculation of Willmore energy (WE), a property related to the Helfrich energy, which, similarly to sphericity, describes the complexity of a shape as defined as its dissimilarity from a sphere (Mondino & Scharrer, 2020; Yoon et al., 2019) (see Materials & Methods for full definition). We find that WE density (Willmore energy weighted by surface area) increases drastically with vacuole complexity, successfully distinguishing between shapes of similar sphericity. While variance of *H* and *K* and Willmore energy can all be considered reasonable metrics for overall shape quantification and comparison, with greater comparative sensitivity than sphericity, Willmore energy density is simpler to interpret. For instance, by mean and Gaussian curvature variances, vacuole I appears to be more complex than vacuole F, however Willmore energy density, which takes both mean and Gaussian curvatures into account, shows that vacuole F is more complex (less sphere-like) than vacuole I. Overall, our curvature analysis demonstrates that the shape information provided by SXT meshes is more precise and realistic than the equivalent measured from voxel reconstructions.

### Examples of detailed features revealed in SXT meshes

The resolution and isotropy of our SXT data, in combination with mesh analysis, reveal detailed features of cells and organelles that are difficult to visualize otherwise, such as biologically relevant surface features, and the spatial relationships between organelles with a high level of detail. We provide several examples of such features here. The vacuole-nucleus junction (NVJ), is a metabolically sensitive signaling platform whose area fluctuates in response to starvation and other functional states (Hariri et al., 2018; Pan et al., 2000). We demonstrate the detection and shape characterization of the vacuole-nucleus interface region, which can be delineated using the known inter-membrane distance range of ∼15-30nm (Hariri et al., 2018), which is comparable to the spatial resolution of our data (Fig. 6A). While the distributions of mean and Gaussian curvatures are visibly distinct when compared across the entire vacuole and nucleus (6A.i. & iii.), the histograms limited to the interface region overlap (6A.ii. & iv.), suggesting that in those regions, the two surfaces match their curvatures at around 0, representing a flat interface. Fig. 6B demonstrates the visibility of minute features in highly complex vacuole clusters, including small indentations, narrow constrictions, and deeply-situated internal holes. The positioning of lipid droplets situated at the periphery of the NVJ or nestled between the vacuole and nucleus (Fig. 6C, top and bottom) are revealed. Mean curvature maps show that the sites of LDs correspond with indentations on the vacuole surface but not on the nucleus. Finally, cell surface geometry visualized by the mesh also reveals the location and number of bud scars, providing a physical landmark as well as potential marker of replicative age for single cells (Fig. 6D). Our approach promises to reveal many more features of biological interest and enable future novel morphological analyses with high spatial precision and statistical throughput.

## Discussion

We have demonstrated that SXT is unique among volumetric imaging modalities in its combination of label-free segmentation, isotropic nanoscale resolution, and capability for statistical throughput (Do et al., 2015; A. Ekman et al., 2020; Guo A et al., 2022; Loconte et al., 2023; Weinhardt et al., 2019). Historically, only optical methods, limited to ∼200 nm resolution (∼100 nm with super-resolution), offered statistically powerful sample sizes (hundreds-thousands), while higher resolution methods (EM, SXT) have been limited in throughput, especially of large volumes such as entire cells, due to laborious manual segmentation. However, the present work and other recent works (Egebjerg et al., 2023; A. Ekman et al., 2020) have demonstrated the capacity for SXT analysis to reach into sample sizes of hundreds of cells, enabling novel quantitative analyses into cellular structure. Here, we have presented a comprehensive approach to studying the subcellular structures of yeast by combining soft X-ray tomography, deep-learning based segmentation, statistical 3D morphometrics, and mesh surface analysis. These findings are significant as they not only provide a clear, quantitative characterization of subcellular components but also highlight the precision and utility of SXT in biological research. Our results highlight the advantages of SXT for generating high quality whole-cell, multi-organelle volumes across large sample sizes, enabling detailed comparisons between individual cells and across groups. The capability to perform statistically-powered population characterization and hypothesis testing with SXT data means that questions previously only accessible by optical microscopy can now be addressed with greater resolution and structural detail, such as comparative analysis of bulk morphometrics across populations.

In the context of disease modeling, the ability to accurately measure and analyze the changes that link genetic modifications directly to phenotypic expressions at the organelle level is essential. The detailed segmentation and subsequent quantitative analysis made possible by our approach have validated its effectiveness in providing crucial insight into how the 3D morphological properties of cellular structures shift in subtle ways across perturbative conditions. We found that our control fluorescence strain, VPH1-GFP, exhibited slight morphological differences compared to the untagged parental strain, which were important to be aware of as a baseline for comparison with *Δvac14*. This suggests that many standardly used fluorescent protein fusion markers and strains which are used for visualization and as controls for perturbative conditions may themselves create subtle off-target alterations in cell structure or function. Our statistical characterization of *Δvac14* also shows that the morphological consequences of the mutant extend beyond reported characterizations such as “gross enlargement” and impaired fission of the vacuole (Bonangelino et al., 1997; Jin et al., 2008; Rudge et al., 2004) to a more nuanced vacuole phenotype in addition to whole-cell effects. This is an example of the fact that morphological phenotypes of yeast mutants are likely far more complex than is reflected in the original qualitative characterizations from low imaging resolution in genetic screens and in concise summaries in Saccharomyces Genome Database (Cherry et al., 1998). With the combined spatial and statistical resolution we report here, we demonstrate the potential to more thoroughly characterize and compare morphological profiles of cell types and strains than previously possible.

The unique combination of data resolution, quality, and throughput achievable through SXT auto-segmentation also offers to open many novel avenues of research. SXT overcomes limitations of other tomography methods such as fixation artifacts, missing wedge artifacts and open edges thanks to a complete 180-degree tilt series and a capacity to image and reconstruct entire cellular volumes, with resolution on the scale of small sub-organelle features (tens of nanometers). SXT is therefore ideally suited for surface modeling using meshing, unlocking a new avenue for precise shape analysis at scale, including surface-wide mapping and statistical characterization of curvature. While we have shown here only a handful of examples of meshes (Fig. 5 & 6), the ability to first perform voxel-wise morphometrics across our entire dataset enabled us to choose statistically representative cells and pinpoint cells of interest to focus on for deeper shape analysis. In the near future, high-throughput mesh processing and analysis of hundreds or thousands of auto-segmented SXT volumes with existing software such as GAMer 2 (Lee, et al., 2020b) or PyCurv (Salfer et al., 2020) will unlock the full potential of high 3D resolution shape information with statistical power, enabling high-throughput comparative whole-cell analyses of detailed morphometrics and precise shape features of organelles and their interactions. We have described several global shape metrics that can be extracted from voxel-wise and meshed volumes, including sphericity, mean curvatures, and Willmore energy density. The exact metric can be chosen as appropriate for the given question, but ultimately such a concise shape descriptor, measured across hundreds or thousands of cells or organelles, can potentially uncover unprecedented insights into shape variability and complex features of mesoscale structures with statistical and comparative power. Such an approach promises to yield a more nuanced understanding of morphological regulation of organelles within cells, across cell populations, and between cellular contexts and genetic backgrounds. This will lead to new insights into the 3D shape, and, as statistical variation reflects dynamic range, shape dynamics of other organelles, as well as of organelle interactions and organelle networks as whole entities. The highly variable nature of 3D subcellular morphology, owing to a complex interplay of stochastic biophysical dynamics and multifaceted molecular regulatory mechanisms, will require such a statistical view to reveal meaningful new levels of understanding.

**Fig 5.**
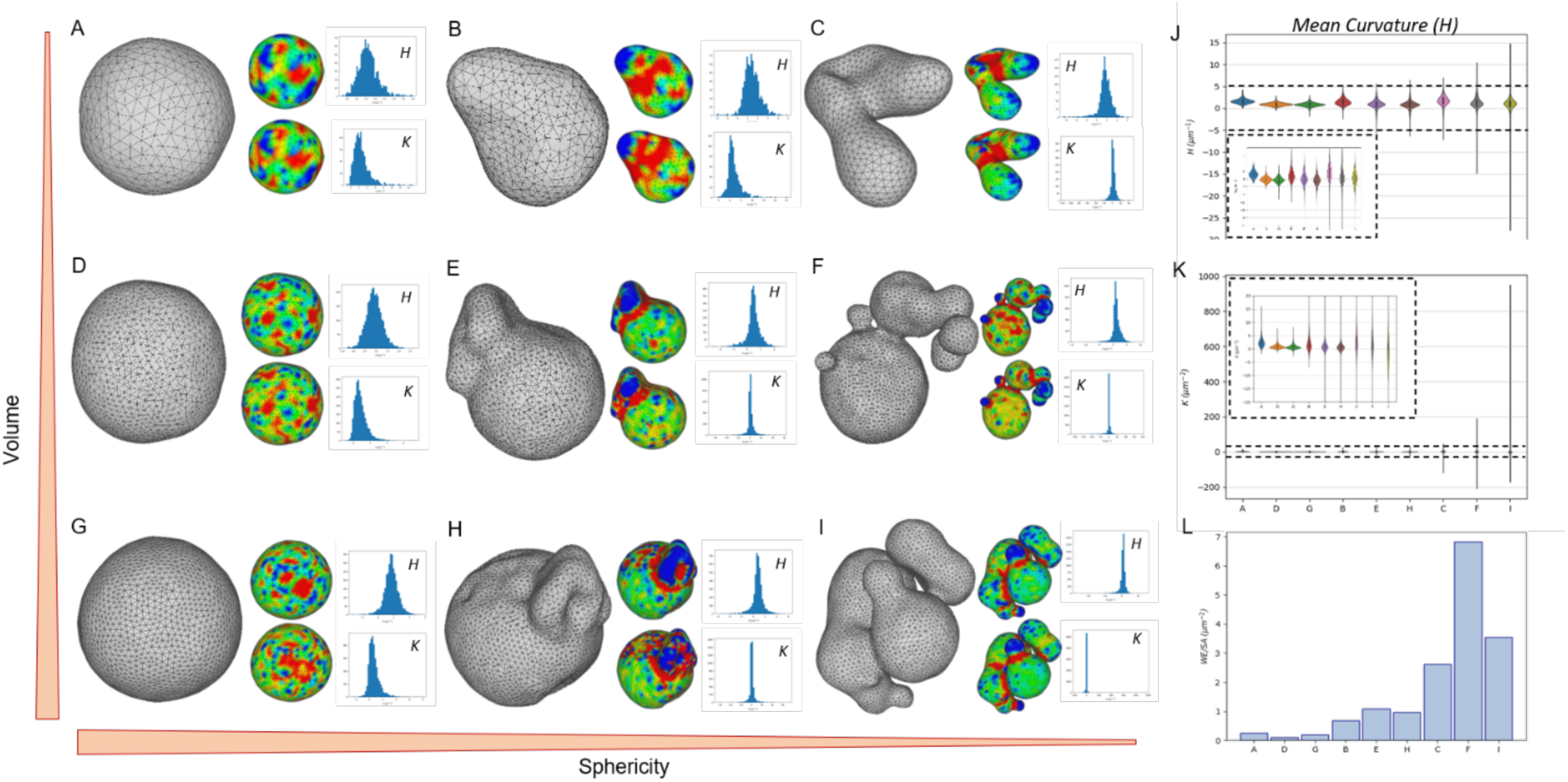
Organelle shape analysis with meshing and curvature mapping. A-I) Example vacuole meshes presented according to increasing shape complexity (decreasing sphericity, left to right, ADG, BEH, CFI) and increasing volume (top to bottom, ABC, DEF, GHI). Curvatures are displayed for each as Mean (*H*) and Gaussian (*K*) curvature maps and histograms (top and bottom, respectively) J-K). Mean (*H*) and Gaussian (*K*) curvature distributions plotted, grouped by volume class and ordered by shape complexity class. Main figure shows full distribution while insets show y-axis zoom. L) Willmore energy density (WE/SA) is calculated as described in Materials & Methods.

The ability to bridge biological scales by detecting fine details of organelles in a whole-cell context unlocks novel multiscale systems-level questions. What are the relationships and interdependencies between fine shape features of multiple organelles? Inter-organelle membrane contact sites (MCS), which are surface regions spanning up to several microns in length (Hariri et al., 2018; Uwizeye et al., 2021a) defined as heterotypic organelle membranes apposed at approximately 10-40 nm distances bridged by tethering protein complexes (Eisenberg-Bord et al., 2016), are challenging to fully resolve by light microscopy. Organelle contacts have been quantified using a range of modalities and resolutions across many cellular contexts, but all approaches are subject to a tradeoff between quantitative precision, volumetric scope, and throughput. For example, it is possible to count organelle associations at the whole-cell scale from voxel co-localization in standard fluorescence data (Shai et al., 2018; Valm et al., 2017; Viana et al., 2023), while electron microscopy enables accurate surface area and distance measurements within limited volumes (Crabtree et al., 2024; Marsh et al., 2001), while FIB-SEM, serial sectioning TEM, and cryo-electron tomography can achieve both, but in small sample sizes (Friedman et al., 2011; Laundon et al., 2019; Uwizeye C et al., 2021a; Uwizeye et al., 2021b). Our work has made strides towards alleviating this tradeoff.

New questions surrounding global intracellular geometric patterns and organelle network arrangements become accessible with the ability to characterize cell-wide changes across cell states and perturbations with high quantitative power. What is the relationship between fine features of organelles and mesoscale cell geometry? How do organelles typically arrange in 3D space? What are the precise changes in organelle morphology, interactions, and intra-cellular distribution involved in complex subcellular remodeling events such as cell division, differentiation, or stress response? How are these changes linked? High-granularity cell and organelle geometries can furthermore be used as basis for structural and biophysical computational modeling, aided by better quality meshes thanks to isotropy (Lee, et al., 2020b; Zhu et al., 2022) as well as mass density information provided by LAC. By quantifying the LAC of various organelles, and relating LAC to other cellular properties, we can gain a deeper understanding of the intrinsic biophysical properties of cells, such as density and compositional variations, which are often influenced by genetic background and cellular health. Integrating molecular, genetic, and dynamic information with the structural constraints and biophysical properties provided by SXT offers a promising avenue for holistic whole-cell modeling capable of predicting complex cell behavior (Loconte et al., 2023). By bridging the sub-micron scale of subcellular architecture with the whole mesoscale anatomy of cells, high-throughput SXT occupies a valuable niche alongside generative 3D modeling (Donovan-Maiye et al., 2022; Majarian et al., 2019; Murphy, 2012), systems organelle dynamics (Mukherji & O’Shea, 2014; Wang & Mukherji, 2022), spatial proteomics (Cho et al., 2021; Hein et al., 2023; Lundberg & Borner, 2019; Qin et al., 2021), morphological profiling (Chandrasekaran et al., 2024; Negishi et al., 2009; Tegtmeyer et al., 2024), and other research areas in the landscape of multiscale integrative cell biology, which looks toward a future holistic understanding of cellular structure and behavior ranging from molecular foundations to complex behaviors within a range of environmental contexts (Goodsell et al., 2020; Rafelski & Theriot, 2024).

## Materials & Methods

### Yeast strains and cell culture

BY4741A (haploid, mating type a), BY4741A *vph1-GFP::his3*, and BY4741A *vph1-GFP::his3 Δvac14* cells were obtained from Jennifer Fung, UCSF. BY4741A and VPH1-GFP::HIS3 have been previously described (Chan et al., 2016; Chan & Marshall, 2014). The *Δvac14* strain was generated by crossing the mutant from the yeast deletion library (Huh et al., 2003) with the VPH1-GFP strain. Cells were grown from frozen glycerol stocks via streaking for single colonies on YPAD agar plates with 30°C incubation. For experiments, 5 mL liquid cultures were seeded with a single plate colony and grown overnight. Fresh cultures were made using a [1:50] dilution of the stationary culture. OD600 was recorded immediately using a tabletop spectrophotometer and after 4 hours of shaking or rotating incubation at 30°C. Log phase cultures (OD600 0.2-0.4) were used for confocal and SXT imaging.

### Confocal imaging and image processing

#### Live organelle staining

1mL aliquots of log-phase cell suspension were incubated with 4 μg/mL DAPI and 200 nM MitoTracker Red FM (ThermoFisher # M7513) at 30°C for 30 min with shaking, then centrifuged at 5000 rpm for 1 min on a tabletop centrifuge, and the pellet was resuspended in 50-100μL PBS. 5 μL of the cell suspension were deposited on 2% agar pads, covered with a coverslip and sealed along the corners with small dabs of clear nail polish.

#### Confocal microscopy

Slides containing live cells at room temperature were imaged on a Nikon Ti inverted microscope with Yokogawa CSU-22 spinning disk confocal, equipped with an EMCCD camera. Published images were taken using a 100x oil Plan Apo VC 1.4 objective with lateral resolution of 93 nm and axial resolution of 100 nm in the GFP (488 nm), DAPI (405 nm), and MitoTracker Red (561 nm) channels to capture entire live cells in z-stacks.

#### Image processing

Confocal images were deconvolved using Huygens software. An experimental PSF was calculated using 0.1 μm TetraSpeck Microsphere fluorescent beads (<u>LifeTech #T7279</u>). Deconvolution settings are as follows: NA 1.4, RI immersion 1.52, embedding medium water 1.338 (standing in for agar pads), backprojected pinhole radius 250 nm, distance 2.53 μm. The background was manually estimated using a histogram generated in the software using a logarithmic vertical mapping function. Deconvolution was performed using an iterative classic Maximum Likelihood Estimation (CMLE) algorithm, using default values (40 max iterations, S:N 20, quality threshold 0.1, optimized Iteration Mode).

Segmentation was performed individually for each channel.

Cell: The cell boundary at the z-slice in the brightfield focal plane was manually traced and filled. This was computationally translated into a 3D volume using a custom algorithm that rotates the 2D segmentation around a central axis in 3D space, creating an ellipsoidal cell volume.

Vacuole: The deconvolved z-stack was processed with a 3D Gaussian Filter (sigma 1), and Despeckle functions, followed by manual thresholding of the processed GFP signal. Nucleus: The DAPI stain marked both nuclear and mitochondrial DNA. To separate the two organelle signals, Mito-tracker was also applied and imaged in a separate channel, and a custom classifier was trained using the Ilastik (Berg et al., 2019) FIJI plugin. The resulting binary images were visually validated against the original fluorescence images and further cleaned with Erosion/Dilation, Fill Holes, and Despeckle processing functions.

### Soft X-ray Tomography (SXT) and image processing

#### Imaging

Live cells were loaded into thin-walled cylindrical borosilicate glass capillaries (Hilgenberg GmbH, Hilgenberg, GER, Cat. No. 4023088) and then vitrified by rapidly plunging them into liquid-nitrogen-cooled (∼90 K) liquid propane. Frozen specimens were cryo-transported to the soft X-ray microscope (XM-2) in the National Center for X-ray Tomography located at the Advanced Light Source of Lawrence Berkeley National Laboratory (Berkeley, CA). At XM-2, the illumination light source is produced by the 1.3 T bending magnet device and directed onto the condenser by a flat nickel mirror. The beam focused by the condenser was order-sorted by the front pinhole onto the specimen and then magnified onto the CCD detector (ANDOR, model iKon-L DO936N BN9KH, 2048 × 2048 pixels) by another micro zone plate. Projection images were collected at 517 eV using a full-rotation imaging goniometer (Le Gros et al., 2014). During data collection, the cells were maintained in a stream of helium gas that had been cooled to liquid nitrogen temperatures (Le Gros et al., 2005; McDermott et al., 2009). Cooling the specimen allows the collection of projection images while mitigating the effects of exposure to radiation. Each dataset (i.e. 90 projection images spanning a range of 180°) was collected using a Fresnel zone plate-based objective lens with a resolution of 50 nm. Exposure times for each projection image were in the range of 150 – 300 ms. The 2D projection images were normalized by the reference image (without a sample), automatically aligned and reconstructed in 3D volumes using the software package AREC3D (Parkinson et al., 2012). The reconstructions do not require fiducial markers or manual interaction with the software and generate virtually real-time initial 3D results with the final 3D volume available within 5 minutes.

#### Auto-segmentation

We used the model described in Ekman, *et al*. (2020) for automatic segmentation, following a 3D U-Net architecture heavily inspired by nnU-Net (Isensee et al., 2021). The model has a depth of five, starting with an initial filter count of 96 at the input layer. Each layer in the encoder path applies 3D convolutional operations, followed by Instance Normalization and ReLU activation. The input data consists of raw LAC volumes binned by a factor of two and processed in blocks of 96x96x96 voxels. We used the same loss function as in Isensee, *et al*. (2021), that combines multi label Cross-Entropy Loss and Dice Loss.

Given the small size of the training dataset, data augmentation was applied to improve model robustness and generalization. Augmentation techniques included random intensity scaling to simulate variations in imaging conditions, along with elastic transformations to warp the input images (Çiçek et al., 2016).

To ensure thorough evaluation and prevent overfitting, we employed a 3-fold cross-validation scheme. The dataset was evenly split into three parts, with each fold serving as a validation set during one-third of the training runs. Training progress was monitored using the validation metric, with early stopping applied when the validation loss reached its minimum.

After training on all three folds, an ensemble approach was used for final predictions. Whole-cell images were generated by fusing an oversampled grid of block-wise label probabilities, using linear blending on overlapping volumes. The prediction probabilities from the three trained models were averaged voxel-wise to produce the final segmentation map. Our code is publicly accessible at https://github.com/ncxt/SXT_AUTOSEG.

### Processing of segmentations

Single cell binary segmentation images were manually examined, slight corrections to the segmentations were performed to fill in holes and remove non-specific structures, and only high quality, complete reconstructions of unbudded cells were retained in the final dataset.

Segmented individually cropped cells were manually examined against original LAC images. Cells were sorted by cell cycle stage according to morphology (unbudded (early G1), budding without organelles in the bud (G1 - early S phase), and recently divided mother-daughter pair. Only unbudded cells were used for the morphometric analysis described here. Incomplete cells (edges cut off beyond the boundaries of the image) and poor segmentations were removed from the dataset, or, when possible, manually corrected using combinations of Erode, Dilate, and Fill Holes functions in FIJI. After QC, a few cells still had non-specific objects segmented, such as small extra structures in the nucleus channel and tiny groups of pixels in the vacuole or LD channels that did not correspond to real organelles. Following morphometric measurements, these structures were filtered out of the dataset by setting a size threshold on the morphometric data determined for each organelle from the volume histogram for the entire population (0.3 μm^3^ for vacuoles, 0.4 μm^3^ for nuclei, 0.02 μm^3^ for LD).

#### Voxel-based 3D morphometrics measurements and analysis

Segmented images were channel-separated such that each structure (cell, vacuole, nucleus, LD) was stored in a separate strain-specific directory. FIJI 3D Suite (Ollion et al., 2013; Schindelin et al., 2012) and Batch Processing was used to calculate 3D measurements in bulk using 3D Volume and 3D Surface functions. All measurements are provided in Source Data file 1.

##### Mesh generation and processing

3D binary segmentation files were surface reconstructed and exported as STL files in FIJI using the 3D Viewer plugin (Schmid et al., 2010), which were then processed in Blender. Each mesh was scaled by adjusting dimensions to known bounding box dimensions obtained from the FIJI’s 3D Objects Counter plugin (Bolte & Cordelières, 2006). Mesh refinement was performed using the GAMer2 plugin for Blender, BlendGAMer, version 2.0.8 (Lee et al., n.d.; Lee, et al., 2020b), using iterations of smoothing and decimation to eliminate terracing artifacts, create even triangulation, and reduce information load while preserving volume and geometry.

Steps: 15 iterations of smoothing, 10 iterations of decimation (Coarse Dense), threshold 1.5, 15 iterations of smoothing, 10 iterations of decimation (Coarse Dense), threshold 1.0, 10 iterations of smoothing, smooth normals. The processed meshes were exported from Blender and visualized in MeshLab (Cignoni et al., 2008), where a final volume-preserving Taubin smoothing step (Taubin, 1995), curvature mapping and measurement, and 3D visualization were performed. Curvature data were exported as .ply files, converted to .csv, and subsequently used for analysis.

##### Willmore energy calculation

In Figure 6, we characterize the extent of deviation from a spherical shape in terms of the Willmore Energy WE (Yoon et al., 2019), which is defined as

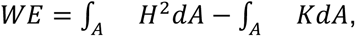

where *H* is the mean curvature, *K* is the Gaussian curvature, and *A* refers to the surface area of the vacuole. MeshLab provides the average and standard deviation of *H* and *K*, averaged over the surface. To calculate the surface integrals from these values, we rewrite the Willmore energy as follows:

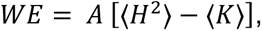

where we use the angle brackets to denote the average. We focus on the Willmore energy density, which is WE divided by the surface area, as previously described (Yoon et al., 2019),

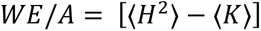

To obtain the first term on the right-hand side, we note that

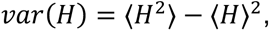

which allows us to compute the Willmore energy density in terms of the outputs from MeshLab as follows:

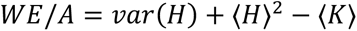

The average values of *H* and *K* are taken over mesh vertices in the software. Here, we convert these vertex-wise averages into integrals over the surface, which is strictly valid only when the mesh face areas are all equal. As a check of this assumption, we found that the integral of the Gaussian curvature calculated over the surfaces in this manner was indeed very close to 4π unless holes were present in the mesh, as required by the Gauss-Bonnet theorem (Pressley, 2010). This assumption will likely be less accurate for more highly convoluted surfaces.

**Fig 6.**
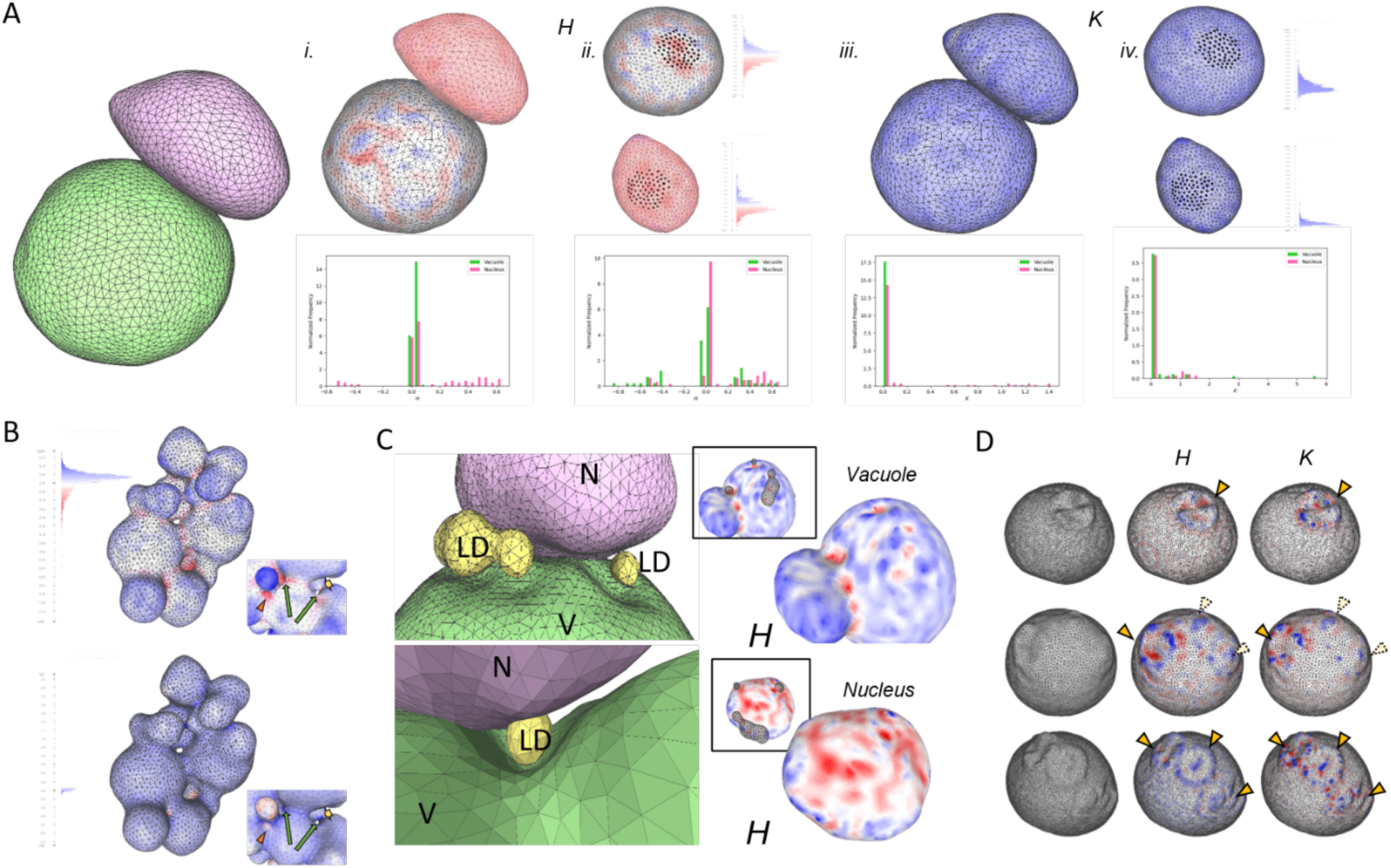
Examples of detailed organelle and cell shape features accessible in SXT data. (A) Contact interface between vacuole (green) and nucleus (pink) i.-ii.) Mean (*H*) and iii.-iv. Gaussian (*K*) curvature shown as colorized surface maps and histograms for whole organelles (i. & iii.) and only the interface region (ii. & iv.). (B) Irregularly shaped vacuole curvature maps reveal detailed surface features. Zoomed views highlight small indentations (orange arrowhead), constrictions between lobes (short yellow arrow) and internal holes (long green arrows). (C) Lipid droplet (yellow) positioning near and between vacuole (green) and nucleus (pink). Mean curvature maps reveal indentations at LD sites (see inset) on vacuole surface (top) but not nucleus (bottom). (D) Bud scars on the cell wall surface. Mean curvature maps aid visualization of one (top), one clear and two putative or newly-forming (middle), and three bud scars (bottom).

##### Vacuole-Nucleus Interface Detection

The Signed Distance Map function in MeshLab was used to compute the signed distance of the vacuole mesh from the nucleus reference, and vice versa, and displayed in a new layer using the Color Mapper tool. Overlapping regions on each organelle were selected using the “Conditional Face Selection” function, setting the face quality threshold as “fq<0”. The selected region was saved in a separate layer and displayed overlaid on the original curvature-mapped mesh as vertices only. Mean and Gaussian curvatures were calculated for only the interface region for each organelle and saved as .ply files for histogram plotting.

#### Data processing, analysis, and visualization

Morphometric and curvature data in .csv files were processed in Python Jupyter notebooks for data analysis and visualization using pandas (versions 1.3-2.2) (McKinney, 2010; The pandas development team, 2020), numpy (version 2.0) (Harris et al., 2020), os, matplotlib (versions 3.5-3.9) (Hunter, 2007), and seaborn (versions 0.12-0.13) (Waskom, 2021). Analysis notebooks are publicly available on Github at https://github.com/mmirvis/SXT-autosegmentation-yeast. 3D rotation movies were taken from MeshLab via screen recording using OBSViewer (version 29.1.3) (*OBS Viewer*, n.d.).

## Acknowledgements

We thank Jennifer Fung (UCSF) and Mark Chan (SFSU) for the gift of yeast strains and advice on fluorescence imaging of organelles in live yeast. Data for this study were acquired at the Center for Advanced Light Microscopy (CALM) Nikon Imaging Center (NIC) at UCSF. We thank Kari Harrington and SoYeon Kim of CALM for training and support with live confocal imaging and deconvolution. We thank Matheus Viana (AICS) for providing the Python script to extrapolate 3D z-stacks for yeast cell images derived from 2D segmentations. We are grateful to Christopher T. Lee and Padmini Rangamani (UCSD) for support implementing GAMer2 and BlendGAMer. We are grateful to members of the Larabell and Marshall groups for helpful discussions and feedback, and to the many investigators whose previous work created the foundation for this work. We acknowledge funding from the Center for Cellular Construction, supported by NSF grant DBI-1548297, and from NIH grant R35 GM130327. Soft X-ray tomography was conducted at the National Center for X-ray Tomography, which is supported by NIH NIGMS (grant no. P30GM138441) and the Department of Energy’s Office of Biological and Environmental Research (grant no. DE-AC02-5CH11231.

## Abbreviations

3D: three-dimensional
CNN: convolutional neural network
EM: electron microscopy
FIJI: FIJI Is Just ImageJ
GAMer: Geometry-preserving Adaptive MeshER software
GFP: green fluorescent protein
LAC: linear absorption coefficient
SXT: soft X-ray tomography
WE: Willmore energy
WED: Willmore energy density

---

**Fig S1.**
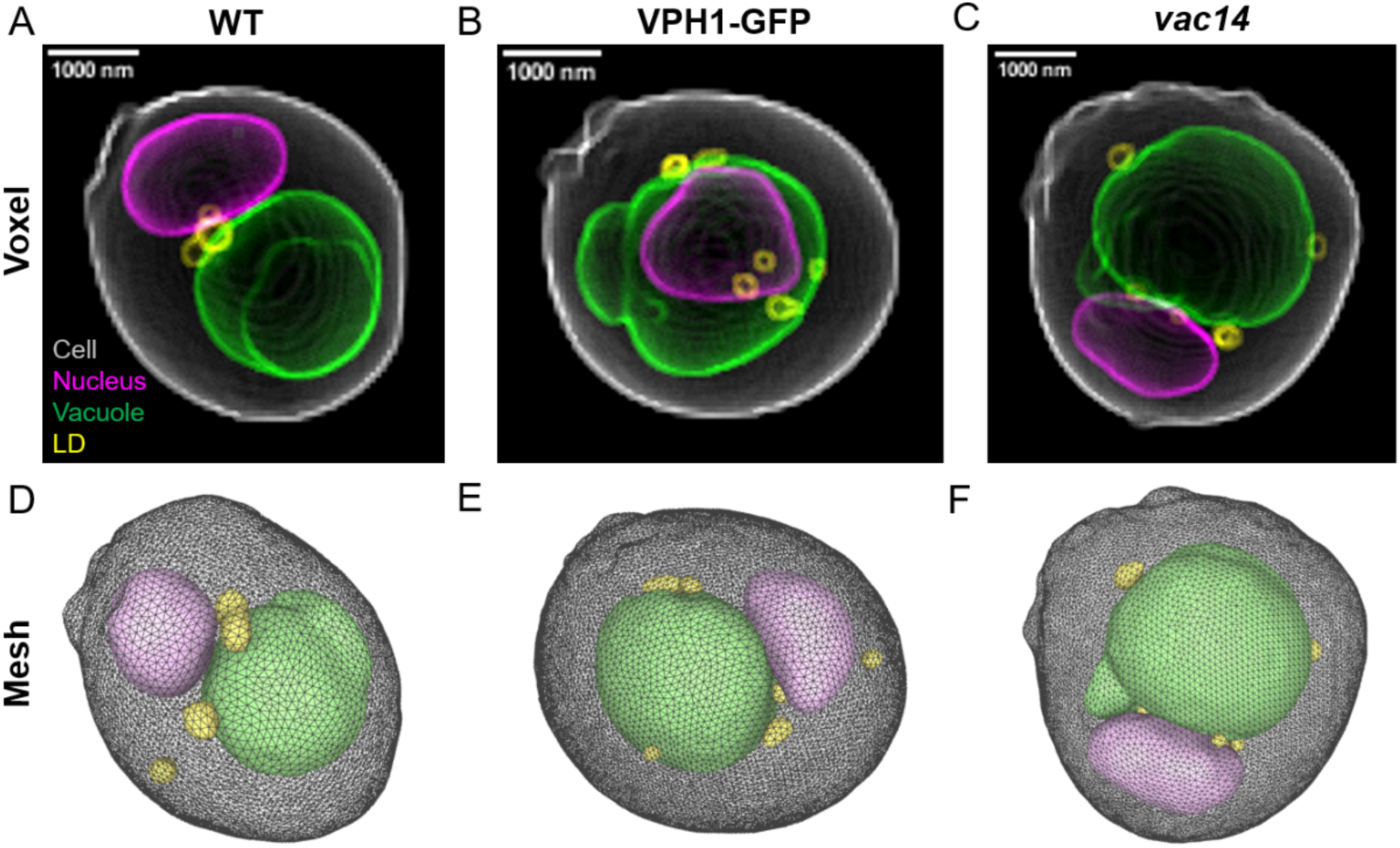
Whole-cell voxel-based and mesh renderings of statistical representatives from each strain. (A-C) Voxel-based reconstructions and (D-F) refined mesh renderings of whole cells including organelles are shown for statistically representative cells from (A,D) WT, (B,E) VPH1-GFP, and (C,F) *vac14* strains, including nucleus (magenta/pink), vacuole (green), lipid droplets (yellow, SXT only), and cell wall (white/gray). See corresponding rotation movies in Supplementary Movies 5-10. Measurements provided in Supplementary Table 2.

